# The influence of 2-aminoethoxydiphenyl borate on the electrical activity of the rat heart under hypoxia/reoxygenation conditions

**DOI:** 10.1101/845909

**Authors:** Elena Kharkovskaya, Grigory V. Osipov, Irina V. Mukhina

## Abstract

The aim of the study was to apply the multi-electrode mapping method to investigate the effect of 2-aminoethoxydiphenyl borate (2-apb) on the electrical activity and vessels of the isolated rat heart under hypoxia/reoxygenation injury (H/R). To date, the ability to influence on the calcium currents and condition of gap junctions has been shown for 2-apb. Changes in the intracellular calcium concentration in the heart play the pivotal role in the development of H/R.

As a result, it was shown that 2-apb influences on the myocardial conduction velocity, sinus rhythm and coronary blood flow in the isolated rat heart. 2-apb caused heart fibrillation during normoxia but not under hypoxia, and attenuated the effect of the H/R on heart rate and vascular function.

## Introduction

2-aminoethoxydiphenyl borate (2-apb) - is the chemical, originally introduced as an inhibitor of Inositol trisphosphate (IP_3_)-induced Ca^2+^ release [1]. Later, it was shown that 2-apb was able to modulate calcium currents without the participation of IP_3_-receptor [2]. In particular, 2-apb can affect store-operated calcium entry (SOCE) – the Ca^2+^ influx across the plasma membrane, which activated in response to depletion of intracellular Ca^2+^ stores in the endoplasmic reticulum [5, 6]. It has been found that 2-apb regulates the ion channels involved in the formation of calcium currents from the Transient receptor potential [5, 6] and Calcium release activated channels [7] families, uncouples gap junctions [8, 9, 10], and other. The 2-apb target depends on the taken concentration and the object of study.

Changes in the cell calcium concentration are important for the heart functioning both in normal conditions and in many physiological disorders, for example, in ischemia/reperfusion injury (IRI). IRI – is the condition in which the resumption of perfusion of the heart after ischemia does not lead to the restoration of its normal functioning, but, conversely, to aggravate the consequences of ischemic effects [11]. This phenomenon is interesting from the point of view of both practical medicine and scientific theory. The effect of IRI is largely due to the effect of hypoxia/reoxygenation (H/R) injury, and the main causes of disturbances in the functioning of the heart and blood vessels are considered to be increased calcium overload and the formation of reactive oxygen species (ROS) [11].

The purpose of the presented research was to study the effect of 2-apb on the myocardial electrical conductivity and the rhythm of contraction of the isolated rat heart with using the multi-electrode mapping method, as well as on the change in coronary blood flow (CBF) under H/R conditions.

## Material and methods

All experimental protocols in this study were approved by the Bioethics Committee of Nizhny Novgorod State Medical Academy, and experiments were carried out according to Act 708n (23.08.2010) of the Russian Federation National Ministry of Public Health, which states the rules of laboratory practice for the care and use of laboratory animals, and the Council Directive 2010/63EU of the European Parliament (September, 22, 2010) on the protection of animals used for scientific purposes.

Wistar rats weighing 200-250 g were anesthetized (Zoletil 100, Virbac Sante Animale, 35 mg/kg, IP), then isolated hearts underwent retrograde perfusion according to the Langendorff method with Krebs-Henseleit solution (sKH): NaCl 118, KCl 4.7, CaCl_2_ 2, MgSO_4_ 1.2, KH_2_PO_4_ 1.2, NaHCO_3_ 20, glucose 10 mmol/L; pH 7.3 – 7.4; at a temperature of 37 C°; under a pressure of 80 mm of water column; saturated with carbogen (95% O_2_ and 5% CO_2_). 2-apb (Sigma, USA) was diluted in sKH (sKH+2-apb), 10 μmol/L. H/R conditions were simulated by perfusion the heart with carbogen-free sKH followed by switching the perfusion to the sKH which was saturated with carbogen. The experimental protocol is presented in table 1.

**Table 1.**
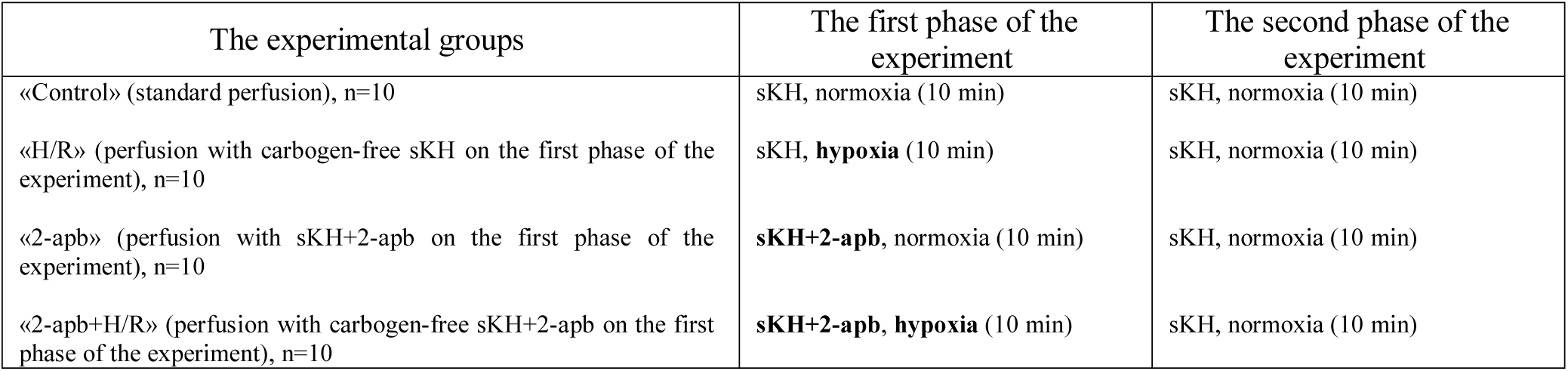
The experimental protocol.

Registration of the electrical activity of the hearts was performed from the surface of the left ventricle with using the flexible multi-electrode arrays MEAFlex72 (Multichannel systems, Germany) (Fig. 1, A, B). The electrical signals recorded from the electrodes were amplified, filtered, and digitized using the corresponding devices: MPA 32I preamplifier, differential filter amplifier FA64I and the data acquisition system with analog-to-digital converter USB-ME 128-System (Multichannel systems, Germany). Recorded electrical signals were visualized and analyzed with help of software applications Cardio2D and Cardio2D^+^ (Multichannel systems, Germany) (Fig. 1, C, D, E).

**Fig. 1.**
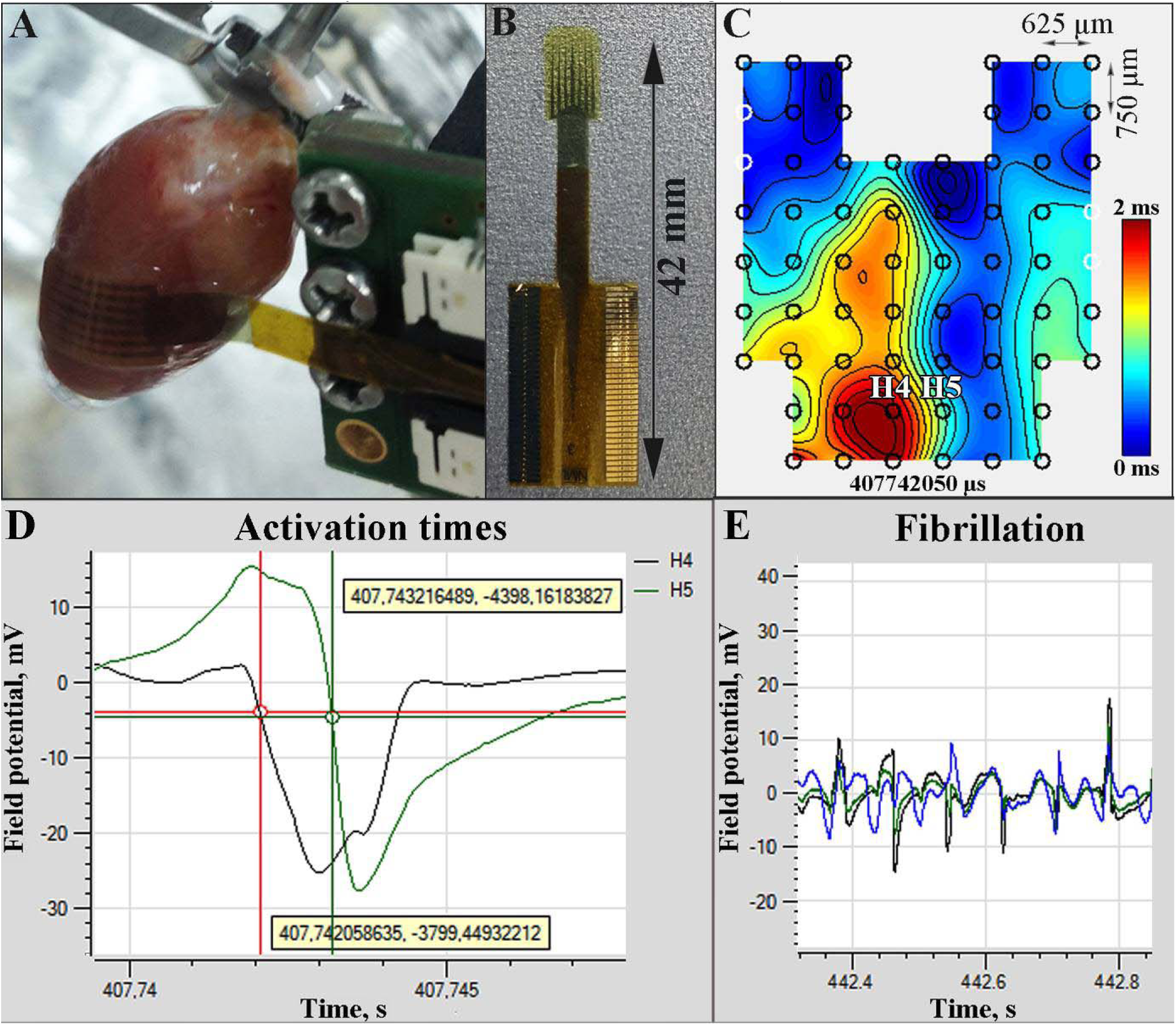
Registration of the electrical activity of the isolated rat heart: **A** - superposition of the multi-electrode array on the surface of the left ventricle; **B** - multi-electrode array MEAFlex72; **C** – isochronal maps (isochrone=100 mV) constructed by the local activation times, which were detected on the electrodes (black circles), electrodes with no local time detected are marked by the white circles; H4 and H5 – is the selected pair of the electrodes for determining LATD; D – the local activation times of the H4 and H5 electrodes corresponding to the maximum downslope of the value of the field potential (y axis) over time (x axis); E - the example of the recording of field potentials from several electrodes during fibrillation

The sufficient amount and configuration of the electrodes in the arrays allowed us to evaluate the value of the local activation times delay (LATD) between two selected electrodes on the epicardial surface. LATD is inversely proportional to the electrical conductivity of the myocardium. The electrodes for assessing LATD were selected based on the isochronal maps constructed in the Cardio2D^+^ program (Fig. 1 C). Myocardium is characterized by anisotropy - different conductivity of the electrical signal depending on the direction (in the longitudinal direction it is higher than in the transverse) [13]. It is generally accepted that anisotropy is due to the elongated shape of cardiomyocytes and the difference in the amount of gap junctions between the lateral surfaces of cardiomyocytes and intercalated discs [14]. In this work the sinus rhythm of the heart was investigated, the electrical stimulation was not applied. That’s why the electrical propagation pattern hadn’t ellipsoid shape because epicardial activity was affected by transmural propagation and Purkinje conduction. But still we could select the one pair of the electrodes with the longest value of LATD within one wavefront, and these values was considered as the value of the transverse conduction of the electrical excitation (Fig. 1, D, H4 and H5 electrodes). By this way, we evaluated the change in the duration of the transverse conduction in the myocardium for each heart, depending on the time and conditions of the experiment. The activation times at the electrodes were determined as the maximum downslope of the value of the field potential. It was problematically to assess the value of LATD during fibrillation because of inability to correctly determine activation times (Fig. 1, E).

Also, heart rate (HR) and coefficient of heart rate variability (CV_HR_) were analyzed as the parameters of the electrical activity of the heart. CV_HR_(%) = 100*standard deviation/average value of interbeat interval (IBI).

The volume of CBF (ml/min) was measured as an indicator of the condition of the blood vessels of the heart.

Changes in the percentage of the values of LATD, HR and CBF were evaluated relative to the last minute of the phase of adaptation of the heart carried out under standard perfusion conditions for 10 minutes before the start of the experiment.

Statistical analysis was performed with GraphPad Prism 6 software (GraphPad Software, USA) using the Shapiro-Wilk test and the Friedman test for repeated measures (p <0.05).

## Results and discussions

### Local activation times delay

#### «Control»

In the control group LATD on the left ventricle surface stayed stable and was equal to 1.2±0.5 msec (Fig. 2, A).

**Fig. 2.**
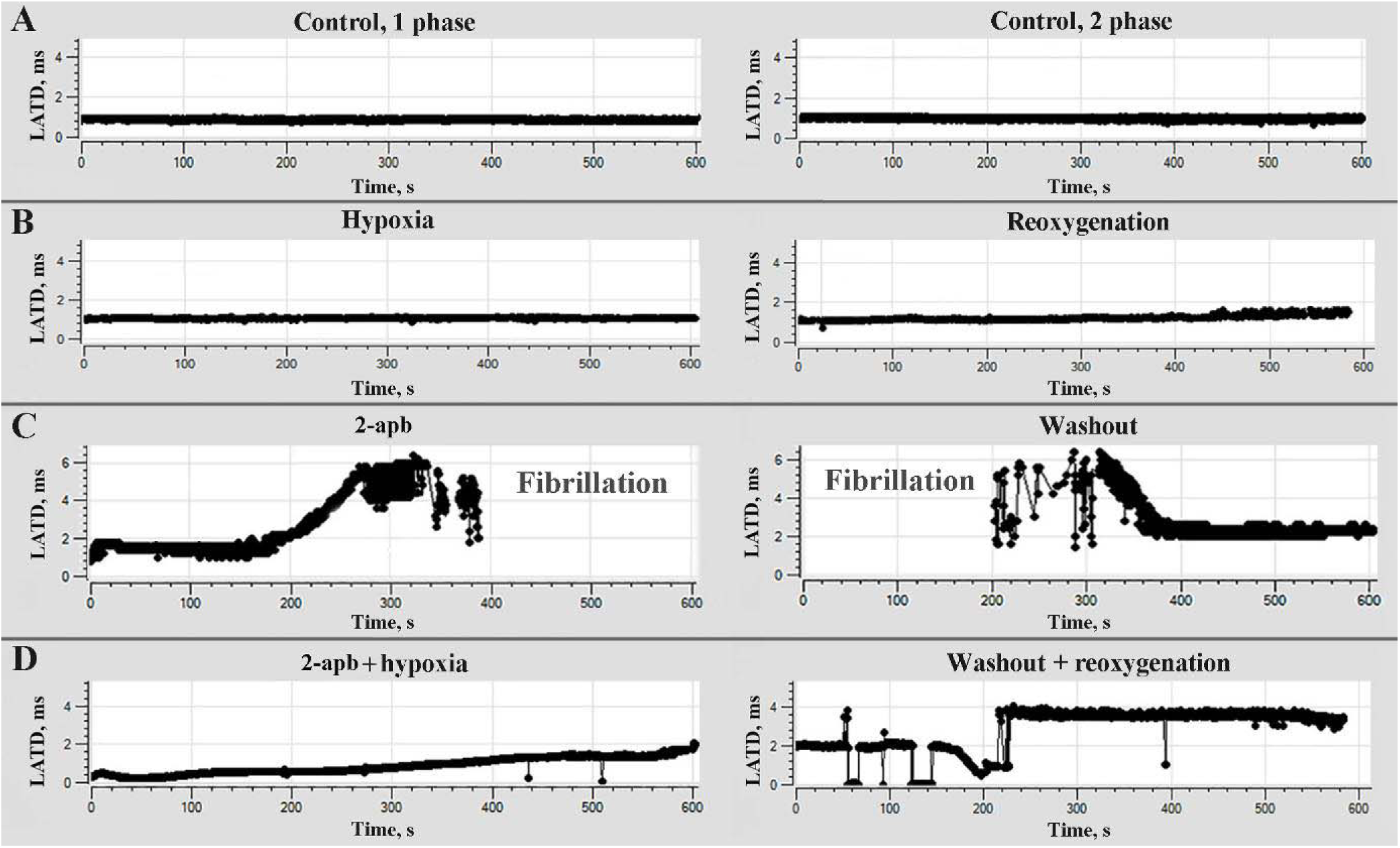
Changes of the values of LATD (y axis) relative to time (x axis) on the examples of the hearts from different experimental groups: **A** – «Control» (n=10), perfusion with standard sKH in normoxia during two experimental phases; **B** – «H/R» (n=10), perfusion with carbogen-free sKH on the first and with carbogen saturated sKH on the second (reoxygenation) experimental phases; **C** - «2-apb» (n=10), perfusion with sKH+2-apb, 10 μmol/L, on the first and with standard sKH on the second (washout) experimental phases; **D** - «2-apb**+**H/R» (n=10), perfusion with carbogen-free sKH+2-apb, 10 μmol/L, on the first and with carbogen saturated standard sKH on the second experimental phases

#### «H/R»

Hypoxia didn’t influence on the LATD values, but towards the end of the reoxygenation phase LATD was increased by 72.8±9.4% (Fig. 2, B). A decrease in electrical conductivity during reperfusion has also been shown in rat cardiomyocytes [15].

#### «2-apb»

After seven to nine minutes of perfusion with sKH+2-apb heart fibrillation was occurred. And two-seven-fold increase in LATD corresponded to the development of this condition (Fig. 2, C). Slowed myocardial conduction velocity is associated with the possibility of occurrence of re-entrant excitation, predisposing to cardiac arrhythmia [16]. The signals from electrodes underwent significant distortion, that’s why mostly it was impossible to identify LATD values between them. On the second phase of the experiment, after switching the perfusion to standard sKH, fibrillation had stopped on average by the fourth minute. Changes in LATD after the period of fibrillation in different hearts were not of a general nature.

Fibrillation of isolated rat hearts caused by using 2-apb (C=22 μmol/L) was also observed by the other researchers [17]. In our work, heart fibrillation was observed with 2-apb in sKH at a concentration of 10 μmol. At a concentration of 10 μmol/L 2-apb was used by other authors, for example, as an inhibitor or blocker of Transient receptor potential canonical 1 (TRPC_1_) channels [5], IP_3_ receptors [18, 19] and SOCE [20]. And this prevented an increase in [Ca^2+^]_in_ in response to different kinds of biochemical effects in cardiomyocytes [5, 18], in vascular smooth muscle cells [20] and fibroblasts [19]. Thus, the cause of fibrillation in our study could be the mechanisms associated with the effect of 2-apb on the concentration of Ca^2+^ ions in heart cells.

For different types of cells, 2-apb has been shown to inhibit or block Cx40, Cx43, Cx45 isoforms of connexins, which are expressed in the heart [10] in concentrations in the range of 1-100 μmol/L [8, 9, 10]. Since the connexin inhibition by 2-apb is caused in a dose-dependent manner, we can assume that in our work, using of 2-apb at a concentration of 10 μmol/L could leads to uncoupling of gap junctions, and this resulted in increase in the values of LATD and fibrillation because of electrical conduction slowing [21].

#### «2-apb+H/R»

In this study, it was shown that 2-apb caused fibrillation under the normoxia, but not under the hypoxia. During perfusion with sKH+2-apb in hypoxic conditions LATD increased by 18.1-310.5% (Fig. 2, D). And after resumption of normal perfusion conditions, there were no differences in the LATD changes (p<0,05).

If 2-apb at a concentration of 10 μmol/L decreases [Ca^2+^]_in_ [5, 18–20], while under hypoxia [Ca^2+^]_in_ increases [12], then this mutually exclusive interaction of 2-apb and the hypoxia effect could prevent the development of fibrillation on the one hand, and changes in H/R observed with hypoxia on the other.

It was shown that connexins are dephosphorylated during hypoxia [22]. This could disrupt their binding to 2-apb and thus prevent fibrillation. According to some reports, 2-apb is able to bind directly to ROS [23], which are formed during hypoxia [12]. Thus, the concentration of the blocker entering the heart could decrease and become insufficient for fibrillation to occurrence.

### Heart rate and heart rate variability

#### «Control»

HR and CV_HR_ did not change during the experiment (p <0.05) and were 246.7±36.8 bpm (Fig. 3, A) and 1.9±1.2% (Fig. 4, A), respectively.

**Fig. 3.**
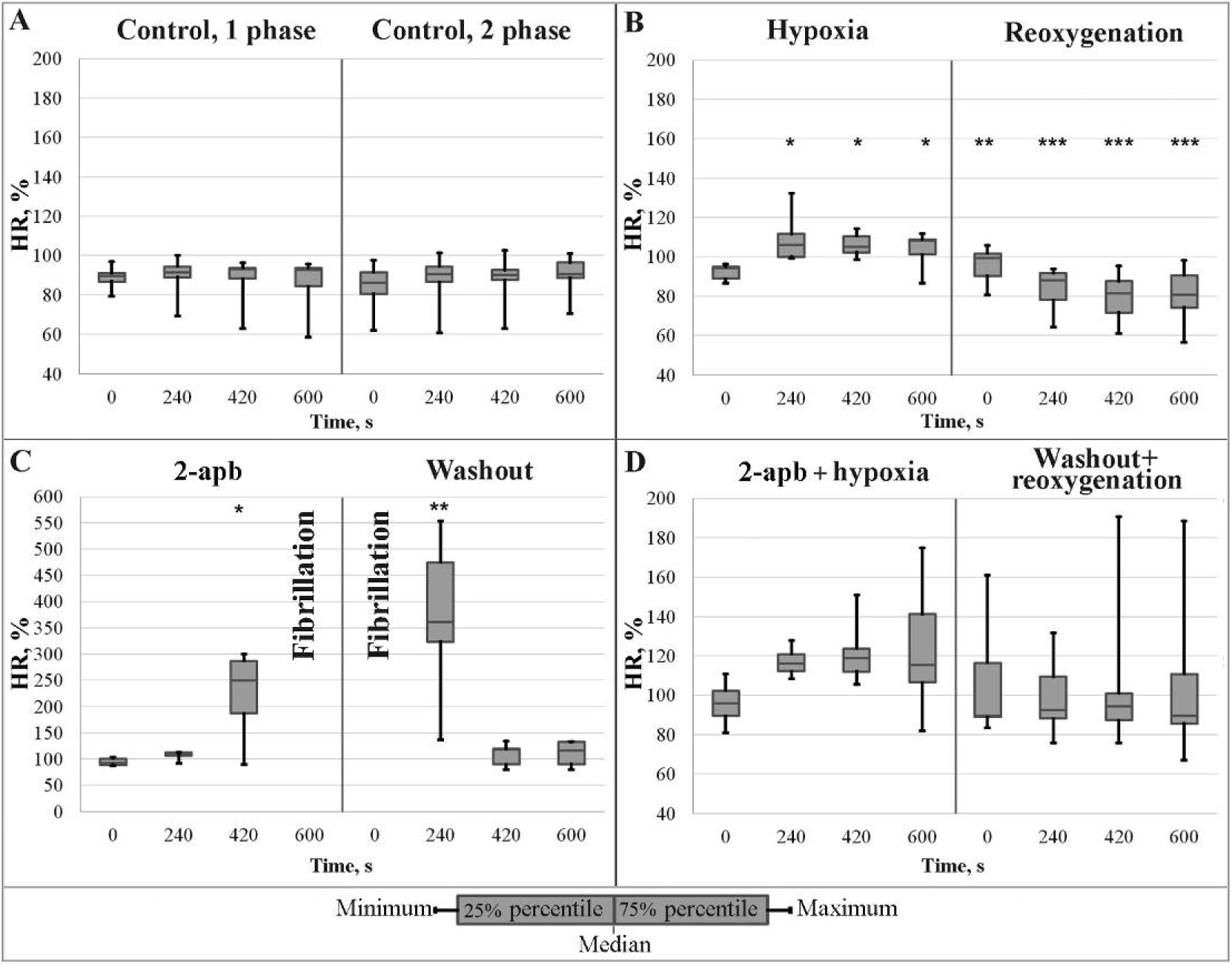
Changes of the values of HR in percentage (y axis) relative to time (x axis) in the experimental groups: **A** – «Control» (n=10), perfusion with standard sKH in normoxia during two experimental phases; **B** – «H/R» (n=10), perfusion with carbogen-free sKH on the first and with carbogen saturated sKH on the second (reoxygenation) experimental phases; **C** - «2-apb» (n=10), perfusion with sKH+2-apb, 10 μmol/L, on the first and with standard sKH on the second (washout) experimental phases; **D** - «2-apb**+**H/R» (n=10), perfusion with carbogen-free sKH+2-apb, 10 μmol/L, on the first and with carbogen saturated standard sKH on the second experimental phases. *, **, *** - the presence of statistically significant differences in the values of HR from the last minute of adaptation (not shown on the graph) taken as 100% according to the Friedman test for repeated measures (p <0.05)

**Fig. 4.**
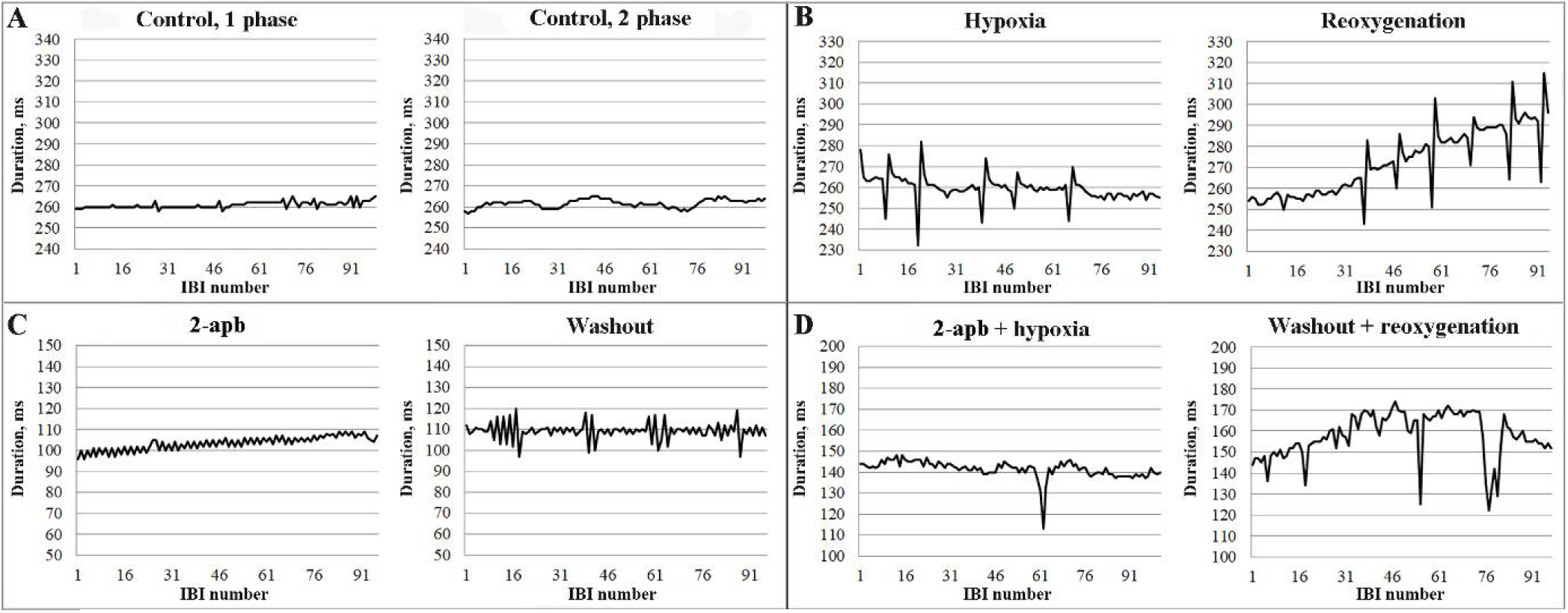
The sequences of IBIs (the values of the time duration of IBIs - y axis; numbers of IBIs – x axis) taken on the third and seventh minutes of the first and second phases of the experiment, respectively on the examples of the hearts from different experimental groups: **A** – «Control» (n=10), perfusion with standard sKH in normoxia during two experimental phases; **B** – «H/R» (n=10), perfusion with carbogen-free sKH on the first and with carbogen saturated sKH on the second (reoxygenation) experimental phases; **C** - «2-apb» (n=10), perfusion with sKH+2-apb, 10 μmol/L, on the first and with standard sKH on the second (washout) experimental phases; **D** - «2-apb**+**H/R» (n=10), perfusion with carbogen-free sKH+2-apb, 10 μmol/L, on the first and with carbogen saturated standard sKH on the second experimental phases

#### «H/R»

Under hypoxia, after 100-200 seconds, HR increased up to 132.4% in some hearts. Whith reoxygenation the fall of HR to 81.3±2.1% was observed (Fig. 2B). During both phases of the experiment rhythm disturbances occurred (Fig. 3, B). Value of CV_HR_ in some hearts reached 12.7%. The lack of oxygen leads to accumulation of Na^+^ and Ca^2+^ ions in the cells due to the inhibition of ATP-dependent transporters, and reoxygenation causes calcium overload [12], which could cause such a picture of HR and CV_HR_ changes.

#### «2-apb»

Significant increase of HR (up to three times) and CV_HR_ (up to 34%) preceded fibrillation (Fig. 3, C; Fig. 4, C). After fibrillation HR stayed increased (370.1±159%) and declined later (Fig. 3, C). CV_HR_ was high and reached up to 14.7% in some hearts after fibrillation on the second phase of the experiment (Fig. 4, C). This could be due to the effect of 2-apb on ionic concentrations in the sinus node pacemaker cells.

#### «2-apb+H/R»

There were no changes in the HR value (p<0.05) in these conditions (Fig. 3, D). CV_HR_ reached a value of 18.33% in some hearts on the first phase of the experiment, and by the end of the second experimental phase ranged from 0.26% to 8.6% (Fig. 3D). It has been demonstrated that pharmacological inhibition of SOCE, which was shown for 2-apb [20], prevents calcium overload at H/ R [24]. And this could offset the effect of hypoxia on HB in these conditions.

### The volume of coronary blood flow

#### «Control»

Under standard conditions of perfusion CBF of isolated hearts gradually decreased on average by 29.3±20.4% by the end of the experiment (Fig. 5, A).

**Fig. 5.**
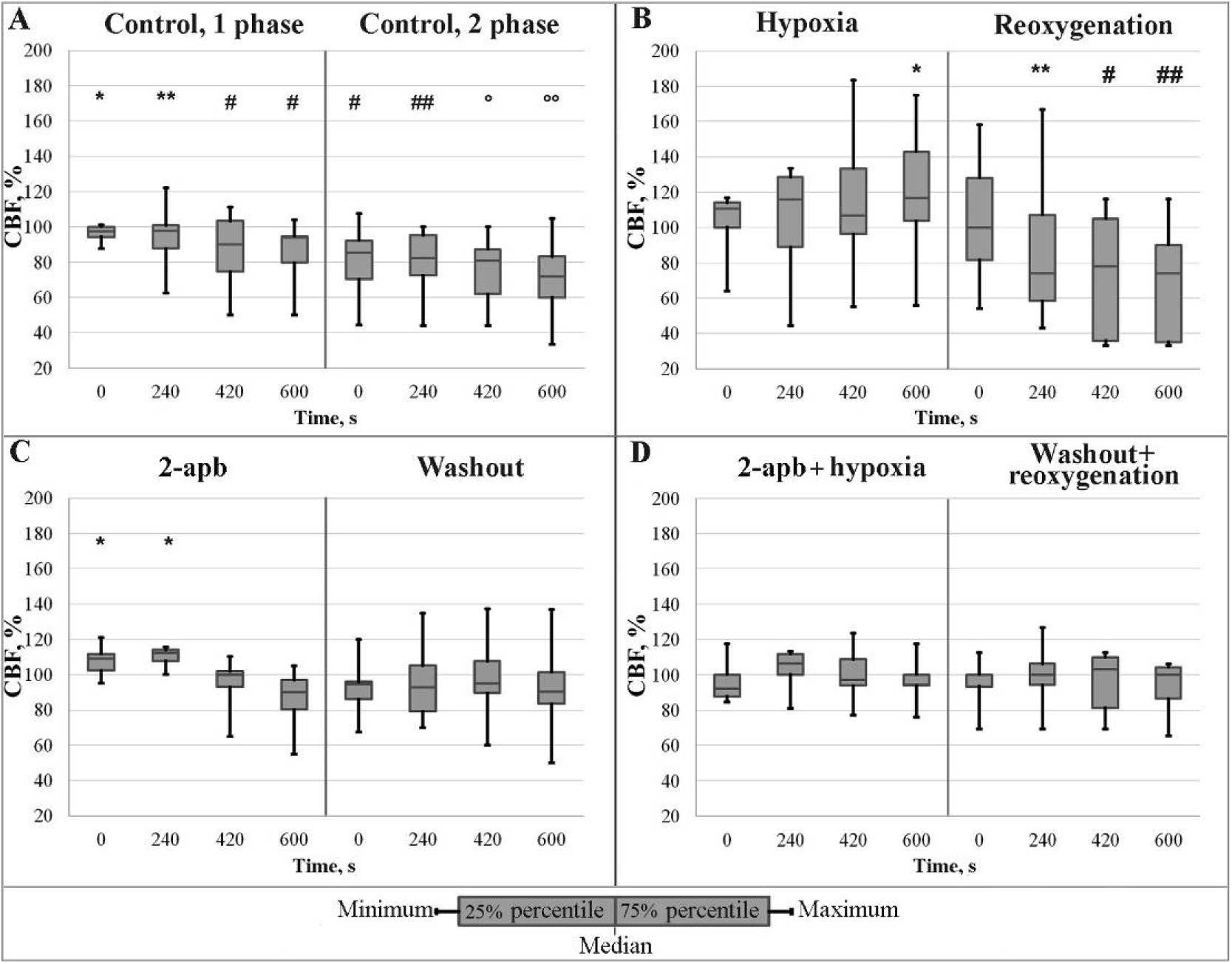
Changes of the values of CBF in percentage (y axis) relative to time (x axis) in the experimental groups: **A** – «Control» (n=10), perfusion with standard sKH in normoxia during two experimental phases; **B** – «H/R» (n=10), perfusion with carbogen-free sKH on the first and with carbogen saturated sKH on the second (reoxygenation) experimental phases; **C** - «2-apb» (n=10), perfusion with sKH+2-apb, 10 μmol/L, on the first and with standard sKH on the second (washout) experimental phases; **D** - «2-apb**+**H/R» (n=10), perfusion with carbogen-free sKH+2-apb, 10 μmol/L, on the first and with carbogen saturated standard sKH on the second experimental phases. *, **, #, ##, °, °° -the presence of statistically significant differences in the values of CBF from the last minute of adaptation (not shown on the graph) taken as 100% according to the Friedman test for repeated measures (p <0.05)

#### «H/R»

Under hypoxia toward to the end of the first experimental phase the value of CBF increased an average by 116.6±40.2%. Presumably, this was due to the vasodilating effect of NO, which is secreted by endothelial cells during hypoxia [19]. Reoxygenation provoked decrease of CBF value to 69.2±32.6% (Fig. 5, B).

#### «2-apb»

In response to the presence of 2-apb CBF was risen during the first five minutes, as maximum of 21%. And then CBF returned to the initial values without changing (p <0.05) until the end of the experiment (Fig. 5, C), in contrast to the decrease in CBF observed under normal conditions of perfusion (Fig. 5, A). The vasodilating effect of 2-apb has been shown by other authors [26]. Disruption of the normal condition of the vessels also can cause the fibrillation [27].

#### «2-apb+H/R»

There were no statistically significant changes of CBF in the in this experimental group (p<0.05). Namely, an increase of CBF caused by hypoxia did not occur in the presence of 2-apb, as well as the CBF decline during reoxygenation, which was observed during 2-apb-free sKH perfusion (Fig. 5, B, D).

## Conclusions

As a result of the study, it was shown that 2-apb affects the parameters of electrical activity and the conditions of the vessels of the isolated rat heart. In particular, perfusion of the heart with a solution containing 2-apb provides:

- the slowing of the myocardial conduction velocity and the development of heart fibrillation, which was observed with normoxia, but not with hypoxia;
- the increase in HR and prevent the decline of this parameter, which was observed under reoxygenation;
- the increase in HR variability, which was more noticeable with in normoxic conditions than under H/R;
- the dilation of coronary vessels and stabilization of CBF, while under hypoxia this parameter markedly increased, and during normoxia and reoxygenation decreased.

The effect of 2-apb can be explained by the influence of this compound on the [Ca^2+^]_in_ of the heart and blood vessels due to the possible regulation of TRPC_1_ channels, IP_3_ receptors, and SOCE [5, 18-20] and by the ability to disconnect gap junctions [8-10]. Changes in [Ca^2+^]_in_ under the action of 2-apb could affect the duration of the action potential of pacemaker cells responsible for the rhythm of heart contractions, which in this paper was evaluated by HR and CV_HR_ parameters. Probably changes in [Ca^2+^]_in_ and uncoupling of gap junctions could influence on the electrical conduction in the myocardium. It can be concluded from the observed decrease in the value of LATD, which was analyzed by the multi-electrode mapping of the heart. The 2-apb-induced changes in [Ca^2+^] in vascular cells were presumably due to the SOCE mechanism and could cause vasodilation, as CBF analysis have shown.

Despite the fact that under normoxia 2-apb causes arrhythmias and fibrillation of the isolated heart, it should be noted its stabilizing effect on CBF under H/R conditions.

For the understanding the mechanism of action of 2-apb on the cardiovascular system, further studies of its properties are required.

## Acknowledgement

This study was financially supported by the Russian Foundation for Basic Research within the framework of the project #17-02-00467.

